# Reconstructing deep-time paleoclimate legacies unveil the demise and turnover of the ancient (boreo)tropical flora

**DOI:** 10.1101/208454

**Authors:** Andrea S. Meseguer, Jorge M. Lobo, Josselin Cornuault, David Beerling, Brad R. Ruhfel, Charles C. Davis, Emmanuelle Jousselin, Isabel Sanmartín

**Author notes:** **Corresponding authors**: Andrea S. Meseguer; Isabel Sanmartín. Real Jardín Botánico, RJB-CSIC, Plaza de Murillo 2, 28014, Madrid, Spain.

## Abstract

**Aim:** Since the Late Cretaceous, the Earth has gone through periods of climate change similar in scale and pace to the warming trend observed today in the Anthropocene. The impact of these ancient climatic events on the evolutionary trajectories of organisms provides clues on the organismal response to climate change, including extinction, migration or persistence. Here, we examine the evolutionary response to climate cooling/warming events of the clusioid families Calophyllaceae, Podostemaceae and Hypericaceae (CPH), and the genus *Hypericum* as test cases.

**Location:** Holarctic.

**Time period:** Late Cretaceous-Cenozoic

**Major taxa studied:** angiosperms

**Methods:** We use paleoclimate simulations, species distribution models and phylogenetic comparative approaches calibrated with fossils.

**Results:** Ancestral CPH lineages could have been distributed in the Holarctic 100 Ma, occupying tropical subhumid assemblages, a finding supported by the fossil record. Expansion to closed-canopy tropical rain forests occurred after 60 Ma, in the Cenozoic, in agreement with earlier ideas of a post-Cretaceous origin of current tropical rain forest. Cooling during this period triggered diversification declines on CPH tropical lineages, and was associated with a climatic shift towards temperate affinities in *Hypericum*. *Hypericum* subsequently migrated to tropical mountains where it encountered different temperate conditions than in the Holarctic.

**Main conclusions:** We hypothesize that most clusioid CPH lineages failed to adapt to temperate regimes during periods of Cenozoic climate change, and thus went extinct in the Holarctic. In contrast, boreotropical descendants including *Hypericum* that underwent niche evolution demonstrated selective advantages as climates became colder. Our results points toward macroevolutionary trajectories involving the altering fates of closely related clades that adapt to periods of global climate change versus those that do not. Moreover, they suggest the hypothesis that potentially many clades, particularly inhabitants of boreotropical floras, were likely extirpated from the Holarctic and persist today (if at all) in more southern tropical locations.

## INTRODUCTION

Climate change is a major agent driving species distribution and diversity patterns (Svenning, Eiserhardt, Normand, Ordonez & Sandel, 2015). Often, studies on biotic impacts of climate change focus on species demographic responses to recent climatic oscillations (Hewitt, 2000), or the effect of current global warming on biodiversity loss (Dawson, Jackson, House, Prentice & Mace, 2011). Yet, short temporal scales might be limited in their capacity to predict the long-term effects of climate changes. The Earth's past climate has gone through periods of climate change that are similar in scale and pace to the warming trend we observe today during the Anthropocene (Zachos, Dickens & Zeebe, 2008). Disentangling the response of organisms to these ancient climate events is key to understand the impact of current warming on biodiversity, but requires the combination of multiple data layers (Svenning et al., 2015), including inferences on the timing of past diversification events, and the paleoclimatic and geographical context under which they occurred. Extinction is likely to have removed important parts of a lineage’s evolutionary history, adding uncertainty. Including fossil information is thus important in deep time ancestral inferences (Slater, Harmon & Alfaro, 2012; Betancur-R, Ortí & Pyron, 2015; Meseguer, Lobo, Ree, Beerling & Sanmartín, 2015). Available paleoclimate data is often restricted to the recent past (Otto-Bliesner, Marshall, Overpeck, Miller & Hu, 2006), limiting as well our ability to infer the impact of ancient events. As a result, studies on deep-time paleoclimate legacies on biodiversity remain scarce (Svenning et al., 2015).

Some of the most dramatic events of climate change occurred during the last ~100 myrs. The Late Cretaceous is considered a relatively warm period. After the mass extinction of the Cretaceous-Tertiary (K/T boundary, 66 Ma) and the impact winter brought by the K/T extinction event, climates became more humid and warmer, with tropical conditions spanning northern and southern latitudes at the beginning of the Cenozoic (Tiffney, 1984; Morley, 2000; Ziegler, Eshel, Rees, Rothfus, Rowley et al., 2003). Global temperatures reached c. 15 degrees warmer than present during the Early Eocene Climate Optimum (EECO), 53–51 Ma (Zachos et al., 2008; Beerling, Fox, Stevenson & Valdes, 2011), but by the Late Eocene (~34 Ma), a precipitous drop in global temperatures termed the Terminal Eocene Event (TEE) (Wolfe, 1985), started a global cooling trend that climaxed during the Pleistocene glaciations. This trend was punctuated with cooling periods such as the Late Miocene Cooling “LMC” (Beerling, Berner, Mackenzie, Harfoot & Pyle, 2009) or periods of warmer climates, such as the Middle Miocene Climate Optimum (MMCO, ~17–15 Ma) or the Mid Pliocene Warming Event (MPWE, 3.6 Ma) (Zachos et al., 2008; Willis & MacDonald, 2011).

The fossil record indicates that these climatic changes were accompanied by concomitant changes in vegetation. The greenhouse interval of the early Cenozoic promoted the extension of tropical assemblages in northern and southern latitudes at the expanse of the subhumid forests of the Late Cretaceous (Tiffney, 1984; Morley, 2000; Ziegler et al., 2003), whereas global cooling at the TEE replaced in the Holarctic a moist, warm-adapted boreotropical vegetation (broad-leaved evergreen and hardwood deciduous taxa) by a temperate mixed-mesophytic forest dominated by coniferous elements (Wolfe, 1975; Tiffney, 1985; Morley, 2007). At the lineage level, paleontological and phylogenetic evidence suggest that the response of plants to these climatic events was diverse, including extinction (Antonelli & Sanmartín, 2011; Xing, Onstein, Carter, Stadler & Linder, 2014; Pokorny, Riina, Mairal, Meseguer, Culshaw et al., 2015); migration (Davis, Bell, Mathews & Donoghue, 2002; Morley, 2007; Couvreur, Pirie, Chatrou, Saunders, Su et al., 2011), or persistence and adaptation, with the evolution of new climatic preferences and relevant traits (Willis & MacDonald, 2011). In the case of the TEE, the latter would have required the evolution of complex physiological systems to cope with frost, such as shedding leaves during freezing periods or senescing above ground tissues (Zanne, Tank, Cornwell, Eastman, Smith et al., 2014). Though there is phylogenetic evidence that adaptation to cold has occurred several times in angiosperms (Donoghue & Edwards, 2014; Spriggs, Clement, Sweeney, Madriñan, Edwards et al., 2015), evidence for the EECO-TEE boreotropical/temperate vegetation transition comes mainly from the paleobotanical literature (Wolfe, 1977; Tiffney, 1985).

The clusioid clade, in the order Malpighiales (Wurdack & Davis, 2009; Xi, Ruhfel, Schaefer, Amorim, Sugumaran et al., 2012), provides an excellent test case to explore the response of flowering plants to rapid climate change, including episodes of global warming (EECO) and cooling (TEE). This clade includes five families: Bonnetiaceae, Calophyllaceae, Clusiaceae, Podostemaceae and Hypericaceae, comprising ~1900 species, and 94 genera (Ruhfel et al., 2011) The families are old, having radiated in a rapid burst in the Late Cretaceous, and considered to be early members of the tropical rain forest (Fig. 1) (Davis, Webb, Wurdack, Jaramillo & Donoghue, 2005; Xi et al., 2012; Ruhfel, Bove, Philbrick & Davis, 2016). Today, clusioid families occur in a variety habitats including closed-canopy rain forest but also open-canopy dry tropical vegetation, temperate and Mediterranean forest, damp and aquatic environments (Stevens, 2007). The majority of clusioid species, however, inhabit the tropical regions of South America, Africa and Southeast Asia, except *Hypericum* (St John's wort, Hypericaceae), the most diverse clusioid genus, c. 500 species, (Robson, 2012), and the only one that has achieved a nearly cosmopolitan distribution across the temperate regions of the world and tropical mountains of the Southern Hemisphere (Meseguer, Aldasoro & Sanmartin, 2013). Previous studies have suggested a link between the evolutionary success of *Hypericum* and the evolution of new temperate affinities relative to their clusioid ancestors (Meseguer et al., 2013; Meseguer et al., 2015; Nürk, Uribe-Convers, Gehrke, Tank & Blattner, 2015). A recent work placed this event of niche evolution in the Oligocene, concomitant with climate cooling and initial diversification in the genus (Nürk et al., 2015). However, phylogenetic and climatic inference in this study did not go further back than the stem-node of *Hypericum* - the divergence between tribe Hypericeae (= *Hypericum*) and tropical sister tribes Vismieae and Cratoxyleae. This limited the ability to explicitly test the niche evolution hypothesis, as information prior to the crown node of Hypericaceae was missing, introducing uncertainty in the inference of ancestral climatic preferences. In addition, fossil evidence suggests that some clusioid lineages were present in the Holarctic before the TEE transition: *i.e.* Late Cretaceous fossil record of Clusiaceae from North America (Crepet & Nixon, 1998) and Paleogene pollen of *Calophyllum* (Callophyllaceae) from Europe (Cavagnetto & Anadón, 1996). At the start of the Mid Cenozoic, however, these families disappear entirely from the fossil record of the northern latitudes. What might explain that disappearance from high latitudes?

**Figure 1.**
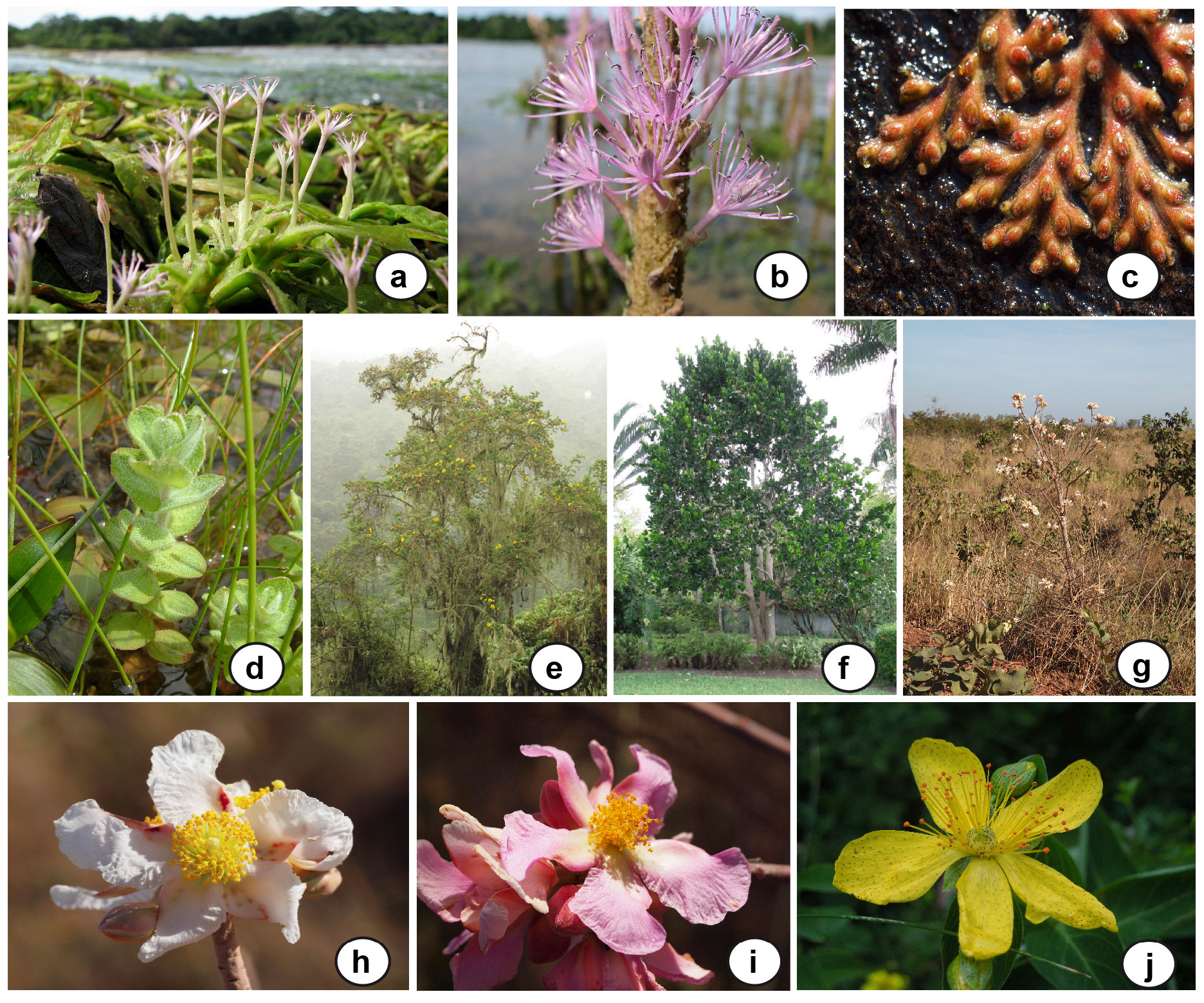
Growth form, morphological, and ecological diversity in the clusioid CPH clade. **(a)** *Apinagia flexuosa*, **(b)** *Mourera fluviatilis*, **(c)** *Castelnavia sp.*, **(d)** *Hypericum elodes*, **(e)** *H. revolutum*, **(f)** *Mammea americana*, **(g)** *Kielmeyera sp.*, **(h)** *Kielmeyera sp.*, **(i)** *Kielmeyera sp.*, **(j)** *Hypericum richeri.* Photographs by B. Ruhfel. *Hypericum* photos by A.S. Meseguer. *H. elodes* photo from www.aphotoflora.com.

Here, we use fossil information in combination with paleoclimate data, species distribution models (SDMs) and macroevolutionary inference methods calibrated with fossils to reconstruct ancestral climatic preferences, past geographical distributions, and diversity dynamics in Hypericum and its closest clusioid relatives: genera within the families Callophylaceae (Podostemaceae, Hypericaceae), herein the clusioid CPH clade. Specifically, our aim is to address the following questions: *(i)* were the ancestors of the clusioid CPH clade adapted to tropical conditions? *(ii)* Is there support for a geographic distribution of CPH lineages in the Holarctic during the Late Cretaceous-Early Cenozoic? *(iii)* When did the modern temperate climatic preferences of *Hypericum* evolved, and have these preferences been conserved over time? *(iv)* Is there a link between the acquisition of temperate affinities and the evolutionary success of *Hypericum* (wider distribution, higher number of species) compared with other clusioid CPH lineages?

## METHODS

### Reconstructing ancestral niche and past geographic distributions

#### Compilation of distributional and climatic data

Extant distributional data for the clusioid CPH clade were obtained from online databases (http://www.gbif.org, accessed on January 2017) and from our own collection. A total of 173.459 data points were collated to exclude ambiguous citations (points in the sea, country centroids, areas out of the native distribution of the species) and repeated occurrences using *SpeciesGeoCoder* (Töpel, Zizka, Calió, Scharn, Silvestro et al., 2017); manual verification was also necessary, as this software did not eliminate all ambiguities. The final dataset included 48.467 occurrences in cells of 5 minutes for Hypericaceae (representing 6 genera and 480 Hypericeae spp. out of 612 described in the family), Podostemaceae (590 occurrences for 38 genera/145 spp. out of 306) and Calophyllaceae (4697 occurrences, 15 genera/276 spp. out of 478) (**Appendix 1** in Supplementary Information). Fossil data (14 records from the Late Eocene to the Oligocene; **Appendix S2**) were obtained from the literature (Meseguer & Sanmartín, 2012; Meseguer et al., 2015). A detailed discussion on the fossil record of *Hypericum* from Meseguer et al. (2015) can be found in **Appendix S3**. Current climatic data were obtained from WorldClim (Hijmans, Cameron, Parra, Jones & Jarvis, 2005) at a resolution of 5 arc-minute cells. For past scenarios we used Hadley-Centre coupled ocean-atmosphere general circulation models, at a resolution of 2.5° × 3.75°, excepting the Cretaceous layer at a resolution 5° × 5° (Beerling et al., 2009; Beerling et al., 2011), which represent six major global warming/cooling events in the Earth's recent history, and designed to incorporate the effects of changing levels of atmospheric CO_2_: 1) Turonian (Cretaceous) (representing the geology and paleotectonics of the time interval 93-89 Ma, and modelled under 1120ppm CO_2_), 2) EECO (53-51 Ma, 1120ppm CO_2_); 3) TEE (34 Ma, 560ppm); 4) MMCO (17-14 Ma, 400ppm); 5) LMC (11.6 Ma, 280ppm); 6) MPWP (3.6 Ma, 560ppm); and 7) the Preindustrial world (> 1900s, 280ppm). The last five were used in Meseguer et al. (2015). The Turnonian paleoclimate model is here used for the first time to accommodate potential climate change during the early history of the clusioid clade. These paleoclimate models include average monthly temperature and precipitation values and were used here to generate seven climatic variables: annual precipitation (AP), annual variation in precipitation (AVP), maximum monthly precipitation (MXMP), minimum monthly precipitation (MNMP), mean annual temperature (MAT), maximum monthly temperature (MXMT) and minimum monthly temperature (MNMT). Among these, we selected the four showing iteratively a Variance Inflation Factor (VIF) lower than 5: AP, MNMP, MNMT and MXMT. VIF quantifies the multicollinearity of predictors (Dormann, Elith, Bacher, Buchmann, Carl et al., 2013), and a value of 5 indicates that the selected predictors are only moderately correlated.

#### Ancestral niche reconstruction

To reconstruct ancestral climatic preferences, we used a composite phylogenetic hypothesis representing relationships at the genus level within the CPH clade and at the section level within *Hypericum* (methods used to construct these trees are described on **Appendix S3**). The clusioid tree was obtained from Ruhfel et al. (2016)’s study and represents 76 of the 94 described clusioid genera, covering all major morphological and biogeographic variation in the group (the first figure in **Appendix S4** as **Fig. S4.1**). The tree was pruned, using *ape* (Paradis, Claude & Strimmer, 2004), to include only representatives of the CPH clade. The *Hypericum* dated tree was obtained from the species-level phylogeny of Meseguer *et al.* (2015), including 114 species and represents all recognized major geographical and morphological variation (**Fig. S4.2**). Because representation of species within sections in this tree was uneven and to avoid bias in the ancestral inference of climatic niches (*i.e.* different number of tips across *Hypericum* sections and clusioid lineages), we pruned the *Hypericum* tree to include one species per major clade recognized by Meseguer et al. (2013) (the first table in **Appendix S5** as **Table S5.1**). Similarly, some genera recovered as non-monophyletic in the CPH clusioid clade (Ruhfel, Bittrich, Bove, Gustafsson, Philbrick et al., 2011) were collapsed to the same branch (see **Table S5.2**). The *Hypericum* tree was grafted into the clusioid tree using the function bind.tree in ape, with a branch length proportional to the age recovered for the crown age of *Hypericum* (Meseguer et al., 2015), which is congruent with the one obtained by (Ruhfel et al., 2016) (**Fig S4.1, S4.2**). The final supertree included 42 tips, representing all major geographic, phylogenetic, and morphological variation in families Calophyllaceae, Podostemaceae and Hypericaceae.

We estimate climatic preferences at the nodes of the clusioid CPH tree using continuous-trait ancestral state inference models (O'Meara, 2012), implemented in the Bayesian software RevBayes (Höhna, Landis, Heath, Boussau, Lartillot et al., 2016), and based on a Brownian model of niche evolution. For each climatic variable, tip values were calculated as the centroid of the climatic conditions of all occurrences assigned to the tip: (max+min)/2. Fossil information was included for *Hypericum* (Late Eocene) crown-ancestors, calculated as the centroid of the climatic condition of Eocene-Oligocene *Hypericum* fossils occurrences in the Late Eocene (560ppm) simulation (**Table S5.3**). The position of this fossil calibration was let as a free parameter along the stem *Hypericum* and integrated upon. We tested the impact of calibration on our estimates by considering all models with and without calibration.

We initially considered more adaptive models, such as the OU or multiple OU models (O'Meara, 2012). However, the selective optimum and the root state cannot be identified under the OU model (Ho & Ané, 2014). We found that this effect concerned all ancestral trait estimates when fitting the OU model on simulated datasets, with highly negatively correlated posterior samples of the selective optimum and ancestral traits. Ho & Ané (2014) further showed that this problem generalizes to multiple-optima OU models. Finally, extensive tests for the fitting of OU models in RevBayes (Cornuault, in prep.) revealed that the inferred number of selective regimes in a multiple-optima OU model, or the uncertainty associated to ancestral trait estimates are highly dependent on the choice of prior for the selection strength parameter of the OU model. In contrast, ancestral traits were generally estimated accurately and with higher precision using a Brownian model, even for data generated under strong OU models (Cornuault, in prep.). Considering these problems for the inference of ancestral traits using OU models, we decided to use the Brownian model.

For the Brownian model, we relaxed the hypothesis of a constant rate of niche evolution by using an uncorrelated exponential relaxed clock for niche traits. Specifically, each branch received its own rate, assumed to be exponentially distributed, with mean *μ*_*σ*_:

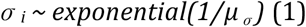

The average rate *μ*_*σ*_ then received an exponential prior of mean 10:

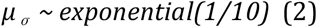

This prior was much flatter than the posterior in all analyses, so it is effectively uninformative relative to the likelihood.

Tip traits were scaled to mean 0 and unit variance before analysis and the root trait received a normal prior with high variance (100) relative to that of tip traits (1):

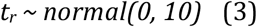

The trait value at the end of branch *i*, *t*_*i*_, received the Brownian motion distribution:

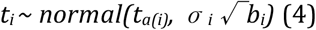

where *b*_*i*_ is the length of branch *i*, and *a(i)* denotes the index of the ancestor of branch *i*.

The time of calibration received a uniform prior between *τ*_*s*_ and *τ*_*e*_, the starting and ending times of the calibration branch (i.e. the stem of *Hypericum*)

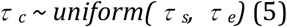

The calibration branch was cut in two parts, with:

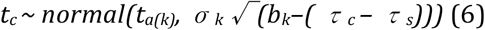

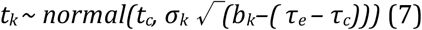

where *k* is the index of the calibration branch and *t*_*c*_ is the trait calibration value (fixed). This model was implemented in RevBayes and three independent runs of 100,000 iterations were carried out with a thinning of 10, providing a final sample of size 30,000 for temperature variables (scripts are available in **Appendix S6**). The posterior distributions of models for precipitation variables were a posteriori conditioned on ancestral traits being positive, by removing all iterations with at least one negative ancestral trait value. This procedure left 29,874 and 19,608 posterior samples for the AP and MNMP variables, respectively. Each MCMC was initially burnt-in for 10,000 iterations, comprising a period of warm-up (wherein operator parameters were tuned) of 5,000 iterations.

To answer the question on whether clusioids were distributed in the Holarctic before the TEE *(ii)*, we represent the potential distribution of major ancestral clusioid CPH lineages, by transferring the reconstructed climatic values of each node (with ASR) to the corresponding paleoclimate simulation. The four climatic variables previously selected and the scale-invariant Mahalanobis Distance (MD) were used to calculate the climatic distance of each cell from the hypothetical “climatic optima” represented by the reconstructed climatic values of each node. The lower quartile of the so obtained MD values was used to represent the most suitable areas.

### Diversification analyses

To investigate the impact of climate cooling on diversification trajectories in the clusioid CPH clade and answer the question on whether a change in climatic preferences triggered rapid diversification in *Hypericum (iv)*, we inferred rates of lineage diversification using stochastic birth-death models and Bayesian MCMC methods in the R package TESS (Höhna, May & Moore, 2016). The supertree described above was used to explore diversification dynamics within the CPH clusioid clade, while for *Hypericum* we used the species-level tree from Meseguer et al. (2015), which allowed us to go deeper into diversification patterns within sections in *Hypericum*. TESS allows comparison among various diversification regimes, including birth-death models where speciation and extinction rates are constant; they vary continuously over time or change episodically in time. The latter (CoMET) correspond to models in which speciation and extinction are allowed to change at discrete intervals, while including also explicit models for mass-extinction (global sampling) events (May, Höhna & Moore, 2016). For the clusioid tree, incomplete taxon sampling was accommodated using the "diversified" strategy in TESS, which is appropriate when sampling is intended to include representatives of major clades, *e.g.* clusioid genera (Höhna, May, et al., 2016). For *Hypericum*, we used uniform random sampling with probability ρ = 0.3. The constant-rate model was specified with two parameters, the speciation and extinction rate, each controlled by an exponential prior distribution with rate = 0.1. We also tested four models with continuously varying diversification rates, in which speciation and extinction rates decrease or increase exponentially over time, while the other parameter remains constant. We used exponential priors for all parameters as above (scripts are available in **Appendix S6**).

Because the number of possible episodic models is infinite, we used reversible-jump MCMC (rjMCMC) approaches in CoMET (TESS) to integrate model uncertainty over all possible episodically-varying birth-death processes including time of rate shifts and mass-extinction events (May et al., 2016). We set the expected number of events (speciation and extinction rate shifts, as well as mass-extinction events) to 2 and used the function *empiricalHyperPriors*=TRUE to determine values of the hyperpriors for the diversification parameters. We run 2 *mcmc* chains for 1 million iterations each, thinning every 100th and discarding the first 1000 as burnin; the auto-tuning function was used in all analyses. We used the package coda (Plummer, Best, Cowles & Vines, 2006) to summarize samples and assess good mixing (ESS) of each *mcmc* run. The Gelman-Rubin test was used to evaluate convergence among runs. Finally, we used stepping-stone simulations to estimate the marginal likelihood of the data under each model (with 1000 iterations and 50 power posteriors), and compare these using Bayes Factors.

## RESULTS

### Ancestral Niche Reconstruction

**Table S5.4** shows extant climatic preferences for clusioid CPH lineages and *Hypericum* clades. **Appendices S7-S14** show for each variable the inferred climatic values with or without fossil calibrations. The ancestor of the CPH clade was found living under similar temperature and precipitation conditions than current tropical non-*Hypericum* Hypericaceae and Podostemaceae taxa (**Fig. 2**; **Fig. S4.3**). The ancestors of Calophyllaceae species thrived under similar temperature but higher precipitation regimes (AP and MNMP) than most other CPH lineages. Stem-node *Hypericum* was estimated to occur in tropical conditions (similar to other non-*Hypericum* Hypericaceae and Podostemaceae taxa), while the ancestral niches within the *Hypericum* clade were characterized by lower temperatures, especially for the MNMT, and precipitation regimes (Fig. 2). The Bayesian posterior distribution of the difference between average trait values for all most recent common ancestors (*mrca*) of *Hypericum* clades (including crown node) and the values estimated for stem-*Hypericum* node yielded 95% highest posterior density (HPD) intervals larger than 0 for the temperature variables, but were non-significant for precipitation variables (**Fig S4.4**). We also calculated the difference between average reconstructed values for all *Hypericum* mrcas and the average *mrcas* of other non-*Hypericum* species in the tree, and found that the HPD for the difference is below 0 for all variables, except for MNMP, which did not differ between the two groups. Finally, we found no difference in trait values between the stem-*Hypericum* node and the average *mrca* of other CPH clusioid taxa for all variables. In sum, our fossil-informed analyses support with confidence that *Hypericum* crown ancestors have on average lower temperature and precipitation (more temperate) values than the rest of the tree. The analyses without fossil constraints produced similar results (**Fig. S4.3**), though differences between average *Hypericum* trait values and stem-*Hypericum* were smaller (**Fig S4.5**) - living under slightly wetter (AP) and warmer regimes (MNMT)- and uncertainty was slightly higher for stem-*Hypericum* reconstructions (Fig 2).

**Fig. 2.**
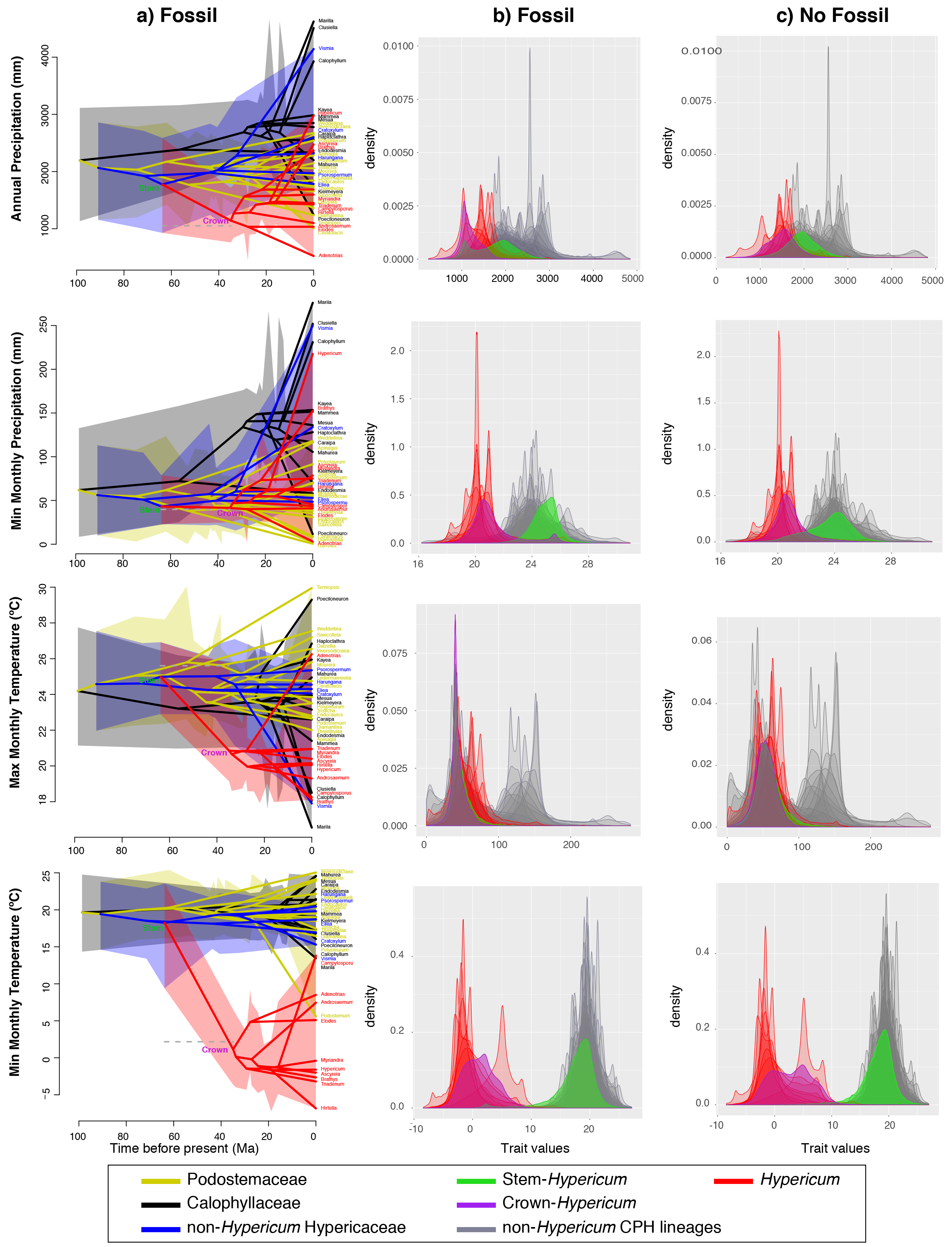
***(a)*** Traitgrams showing the inferred evolution of climatic variables (Annual Precipitation, Min Monthly Precipitation, Min and Max Monthly Temperature), using fossil calibrations and over the dated phylogenetic tree of the clusioid CPH clade in a space defined by the phenotype (y axis). Shaded areas encompass all HPD intervals of the group concerned. The grey dashed horizontal line represents the calibration value. ***(b)*** Posterior distributions of ancestral traits estimated for all nodes using fossil callibrations. ***(c)*** Posterior distributions of ancestral traits estimated for all nodes based on present taxa only.

The geographic projection of these ancestral climatic values onto the corresponding paleoclimate layer (**Fig. 3**) indicates that the ancestors of CPH lineages could have found favourable conditions in the (presently) tropical regions of northern South America and central Africa, the southern coasts of the Northern Hemisphere, and the East Gondwana landmasses of Antarctica, southern South America, and Australia. The geographic extent of these favourable conditions increased in the Holarctic for the ancestors of family Hypericaceae, whereas *Hypericum* shows a continuous presence across the southern parts of the Northern Hemisphere during the entire Cenozoic period, in agreement with projections obtained from fossil-based SDMs (Meseguer et al., 2015).

**Fig. 3.**
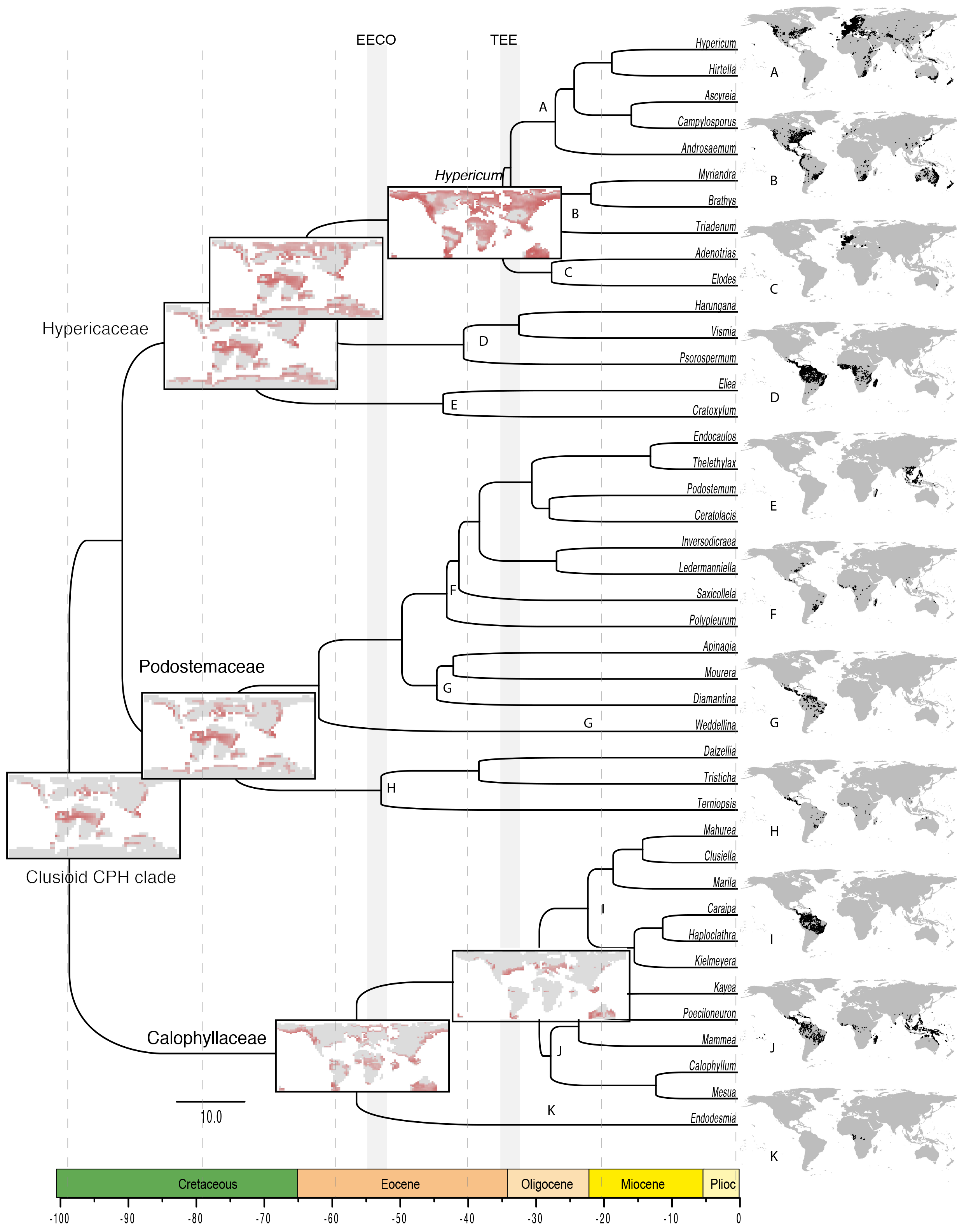
Potential distribution of the clusioid CPH clade in the past (in red) derived from the ancestral state reconstruction (ASR) calibrated with fossils. Estimated values are projected over a suite of paleoclimate reconstructions and plotted over the clusioid CPH clade tree. Maps on the right represent occurrences used in this study and reflect present ranges of the taxa. Abbreviations: EECO = Early Eocene Climate Optimum; TEE = Terminal Eocene Event

### Diversification analyses

For *Hypericum*, the episodic diversification model fitted the data slightly better than the constant rate model (BF > 1.3), but substantially better than other continuous-varying diversification models (BF > 6.7; **Table S5.5**). CoMET (TESS) did not found evidence of mass-extinctions in *Hypericum*, but identified a rate shift (weakly supported) towards increasing speciation and decreasing extinction around 15-10 Ma (BF = 2; **Fig. 4**; **Fig. S4.6**). For the clusioid tree, the constant-rate model fitted the data slightly better than a model with extinction increasing over time (*IncrD*, BF= 1.2) and than the episodic model (BF = 2.2). The latter identified a supported shift (BF > 4) towards decreasing speciation rates during the Late Eocene−Miocene, between 30-10 Ma (**Fig .4**; **Fig .S4.7**). The lack of significant support (BF > 5) for varying diversification rates in our models is likely a result of the small tree size. In any case, for the clusioids, models showing an increase in extinction rates were strongly preferred (BF > 10; **Table S5.6**) over all models with varying speciation and constant extinction.

**Fig. 4.**
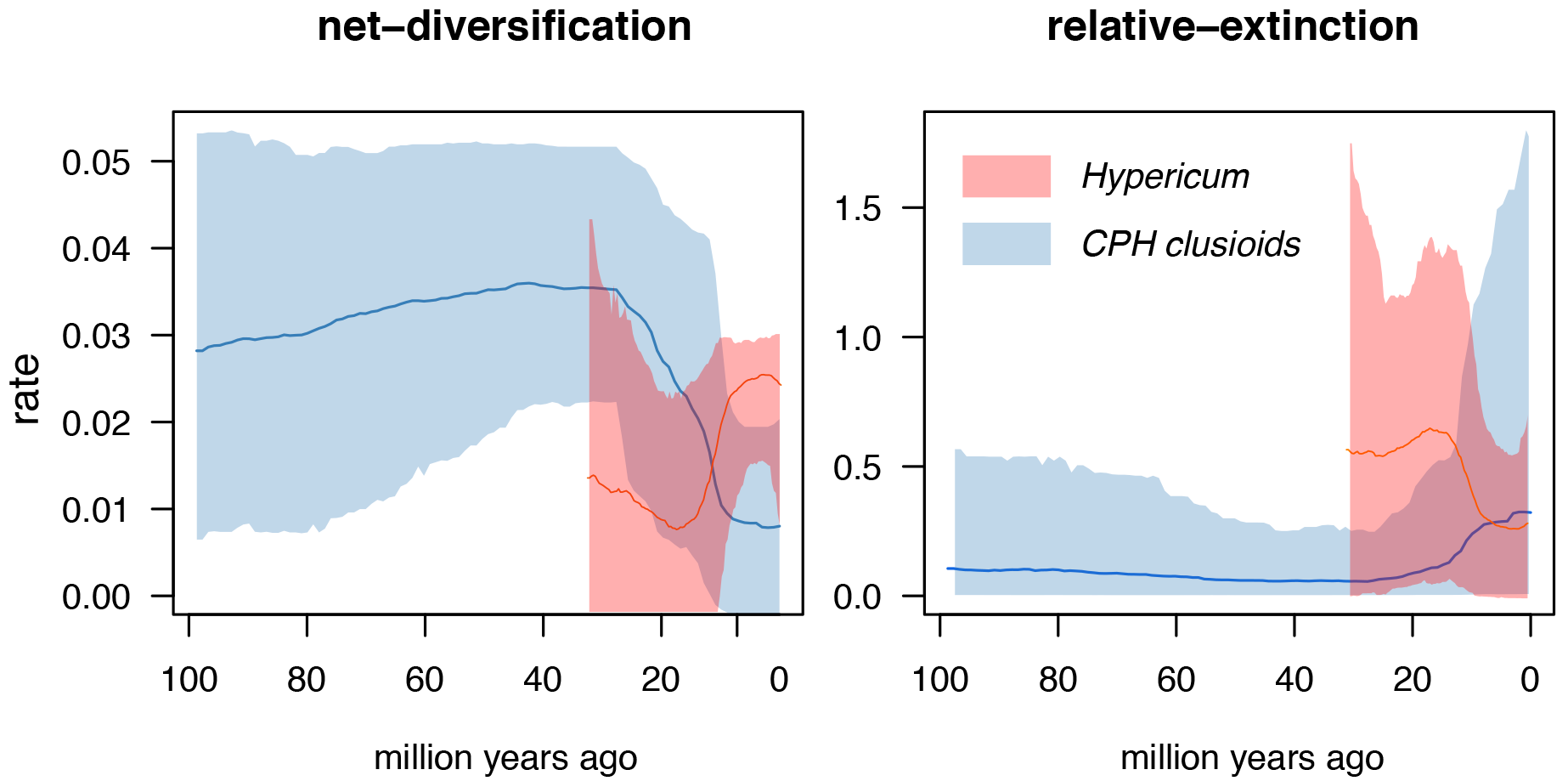
Net diversification (speciation-extinction) and relative extinction (extin-ciont/speciation) of the clusioid CPH clade and *Hypericum* through time as estimated by the episodic model of TESS (CoMET).

## DISCUSSION

### Late Cretaceous-Early Cenozoic evolution of the clusioid CPH clade: boreotropical roots, climate precipitated extinction, and southward migration

Fossil evidence suggests that most Late Cretaceous floras were tropical-like but largely represented open and subhumid forests (Upchurch & Wolfe, 1987). The traditional view is that the current extension of the tropical, closed-canopy rain forest was achieved more recently, following the impact winter precipitated by the K/T mass-extinction (Tiffney, 1984; Morley, 2000; Ziegler et al., 2003). Controversy, however, exists in this regard, with some evidence suggesting that wet-megathermal forest, including angiosperms, arose by the Cenomanian before the K/T (Wolfe & Upchurch, 1987; Davis et al., 2005). Based on current species habitat occupancy, Davis et al. (2005) reconstructed the habitat of the ancestral species of the clusioid clade (~102 Ma; **Fig. S4.1**; Ruhfel et al., 2016) in warm, wet, closed-canopy forests. This reconstruction, however, did not include representatives of family Calophyllaceae, so the habitat occupancy of the CPH clade (~99 Ma) could not be fully inferred. Yet, ancestors of Podostemaceae+Hypericaceae (~91 Ma) were inferred to have lived under drier conditions than the clusioid ancestor, in open-canopy, dry tropical vegetation (Davis et al., 2005). Our paleoclimatic reconstructions based on the inference of ancestral climatic preferences suggest that the Cenomanian clusioid ancestors - in this case the CPH clade - occupied warm subhumid assemblages. Specifically, we inferred that the *mrca* of Calophyllaceae, Podostemaceae and Hypericaceae lived during the Late Cretaceous in regions with similar temperatures but drier precipitation regimes (Annual Prec = 2000 mm) than more recently evolved tropical Calophyllaceae (AP = 4000mm), though similar regions inhabited by most extant Podostemaceae and Hypericaceae species (**Fig. 2**). The evolution of preferences for more humid environments in the CPH clade is suggested to have evolved later during the early Cenozoic (< 60 Ma), and independently in family Calophyllaceae and Hypericaceae. These findings have additional implications for the fossil Paleoclusia chevalieri (90 Ma) from New Jersey (Crepet & Nixon, 1998). As a close relative of the CPH clade - placed within the clusioid clade as stem or crown member of Clusiaceae s.s. (Ruhfel, Stevens & Davis, 2013) - our results raise the possibility that this taxon inhabited warm-subhumid habitats in the Holarctic during the Late Cretaceous as well. This is supported by its northern hemisphere distribution, similar to CPH ancestors (Fig. 3), and by earlier interpretations that this locality was likely characterized by warm subtropical climates (Crepet & Nixon, 1998). Overall, our results are consistent with hypotheses of a post-Cretaceous origin of current tropical rain forest. However, we acknowledge we are considering very large geographical areas that likely encompass potentially high variability in habitats, and fossil evidence from North America suggesting the presence of tropical rain forest before the K/T.

Today, each of the three clusioid CPH families, Callophyllaceae, Podostomeceae, and Hypericaceae, exhibits an amphi-tropical distribution, with different genera inhabiting the Neotropics, South East Asia, and Africa (**Fig. 3**). Ruhfel et al. (2016) argued that this disjunct distribution across former Gondwanan landmasses was achieved by “boreotropical migration” during the Early Cenozoic or long-distance dispersal, rather than by ancient vicariance. The "boreotropical connection" hypothesis posits that current widespread tropical lineages were formerly present in the Northern Hemisphere when climates were warm and humid at the start of the Cenozoic, but became extinct in these regions and moved, or were restricted, to Equatorial regions as climates became cooler following the TEE (Lavin & Luckow, 1993; Davis et al., 2002; Morley, 2007). Our ancestral niche and diversification analyses support this hypothesis. Geographic projections of ancestral climatic preferences show that ancestors of the clusioid CPH lineages could have found favourable conditions in the Northern Hemisphere before Cenozoic climate cooling (**Fig. 3**), which is in agreement with the fossil record (Cavagnetto & Anadón, 1996) and with biogeographic reconstructions supporting the presence of Hypericaceae ancestors (~71 Ma) in the Holarctic (Meseguer et al., 2015). Our continuous diversification analyses also indicate a trend of increasing extinction rates through time in the CPH clusioid clade, while the episodic model supports a scenario with a decline in diversification rates and increase in the extinction fraction towards the end of the Eocene (**Fig. 4**). It is possible that, when the tropical biome retreated to the south (Morley, 2007), these clusioid lineages failed to adapt to new temperate regimes and went extinct, persisting in refugia in southern tropical regions (**Fig. 2**, **3**). Similar scenarios of climate-driven extinction have been inferred in other angiosperms facing climate cooling (Antonelli & Sanmartín, 2011; Willis & MacDonald, 2011; Xing et al., 2014).

### Late Cenozoic evolution of Hypericum: temperate niche shifts, diversification, and geographical expansion into tropical mountains

Meseguer et al (2015) suggested that *Hypericum* stem-lineages were distributed in the Holarctic before the EECO, and could have been part of the Early Cenozoic boreotropical flora. Our results support this hypothesis. Climatic preferences for *Hypericum* stem-node were inferred to be more similar to those observed today among other tropical Hypericaceae taxa - and other CPH lineages - than to extant *Hypericum* species (**Fig. 2**). Geographic projections of these climatic preferences show climatically suitable conditions for stem-*Hypericum* in the Holarctic during the Paleocene (~64 Ma; **Fig. 3**). By the TEE, concurrent with climate cooling (~35 Ma), the *mrca* of all extant species (*Hypericum* crown-node) was already inferred to be adapted to temperate conditions: tolerating average temperatures up to −6°C during the coldest month of the year, and drier environments than their ancestors, with less than 1000 mm of annual precipitation (**Fig. 2**, **Appendix S7-S10**). This supports the idea that an event of climate niche evolution along the stem-branch of *Hypericum*, the acquisition of new temperate tolerances, allowed this genus to persist in the Holarctic after the TEE dramatic transition, whereas other clusioid relatives went extinct (Meseguer et al., 2013; Meseguer et al., 2015). A previous study (Nürk et al., 2015) has also inferred a climatic niche shift towards temperate tolerances at the onset of *Hypericum* diversification, but this event is dated later during the Oligocene. The estimated Oligocene age for crown-*Hypericum* by Nürk et al. (25.8 Ma, 95% HPD=33-20 Ma; cf. Fig. 2) is about 10 Ma younger than the one estimated in Meseguer et al. (2015)-Late Eocene 34.9 Ma (HPD=34-37 Ma, **Fig. S4.2**). It also differs from Ruhfel et al. (2016) analysis of the clusioid clade (37.3 Ma, HPD=26-52, **Fig. S4.1**), which employed a different set of fossils, and from a more comprehensive analysis of Malpighiales dating the crown group of the New World clade of *Hypericum* at 25 Ma (Xi et al., 2012), which is otherwise in agreement with our estimates for this node (29 Ma; HDP 23-33). Nürk et al.'s biogeographic reconstruction is also in conflict with the boreotropical hypothesis (stem lineages were reconstructed during the greenhouse period of the Eocene, including the EECO, in Africa) and in disagreement with the fossil record (Meseguer & Sanmartín, 2012) (see further discussion in **Appendix S3**).

Were these temperate preferences conserved over evolutionary time in *Hypericum*? Overall, temperate preferences similar to those of crown-node *Hypericum* are shared by clades such as North American section Myriandra or semiaquatic Mediterranean Elodes (**Fig. 2**). However, a trend towards increasing tolerance to cold can be observed in some clades, such as the *mrca* of Hirtella, with species widely distributed in Eurasia. Three *Hypericum* lineages were reconstructed in Meseguer et al. (2013) as migrating into the southern tropical mountains after the LMC event (c. 10 Ma): *Campylosporus s.l.* clade in Eastern Africa, *Brathys* in the Andean Paramo, and *Ascyreia* in the Himalayas. Although the common view is that the Miocene-Pliocene uplift of these mountains opened new areas with similar climatic conditions for temperate taxa, which promoted rapid diversification in pre-adapted taxa (Graham, Carnaval, Cadena, Zamudio, Roberts et al., 2014; Merckx, Hendriks, Beentjes, Mennes, Becking et al., 2015), our results suggest that *Hypericum* did not experience the same temperate conditions in these regions than in the Holarctic. Species of *Ascyreia* and *Brathys* occur in more humid environments than those inhabited by other extant *Hypericum* species (excepting in section *Hypericum s.str*.), and than those inferred for their *Hypericum*-crown ancestors (**Fig. 2**). Their precipitation tolerances are indeed more similar to those of other tropical clusioid CPH lineages. Temperature tolerances for *Campylosporus* are also the most similar to the ones inferred for tropical stem-*Hypericum*. All of this suggests a remarkable ability of *Hypericum* to occupy different environments within the temperate biome, with climatic preferences being to a certain degree phylogenetically labile in this lineage (Evans, Smith, Flynn & Donoghue, 2009; Wüest, Antonelli, Zimmermann & Linder, 2015).

Was the shift to temperate tolerances key for the evolutionary success of *Hypericum*? A general tenet in evolutionary ecology is that the acquisition of new climatic preferences allows a lineage to colonize novel ecological niches (*e.g.* temperate habitats), which can promote rapid diversification by ecological release, *i.e.* the availability of novel resources and reduction in direct competition (Wiens, Ackerly, Allen, Anacker, Buckley et al., 2010). We found no direct correlation between the event of niche evolution before crown-*Hypericum* (> 35 Ma) and an increase in diversification in the genus. Instead, we found that extinction rates decreased and speciation rates increased in *Hypericum* after climates became increasingly colder with the Late Miocene climate cooling, c. 10 Ma (**Fig. 4**). This increase in diversification in the Late Miocene coincides with Meseguer *et al.* (2013)'s reconstruction of migration events into the newly uplifted tropical mountains during this period within *Brathys* (Andes), *Ascyreia* (Himalayas), and *Campylosporus* (Africa), and with the inference of potential climatic adaptation in these clades (**Fig. 2**). In sum, rather than the acquisition of temperate affinities at the start of *Hypericum* evolution promoting rapid diversification, the extant high diversity levels observed today can be explained by a combination of: *1)* geographic range expansion facilitated by plate tectonics - *i.e*. the Miocene connection between North and South America, or between Europe and Africa - and Late Miocene climate cooling creating novel temperate niches in tropical mountain regions, and *2)* niche evolution to new temperate wetter conditions (*e.g. Brathys, Ascyreia*).

### Niche evolution, climate change and the boreotropical roots of temperate taxa

The TEE was probably the most dramatic climate cooling of the last 66 Ma, representing a vegetation turnover at a scale (temporal, geographic, compositional) not seen later (Morley, 2007), and has often been associated to historically high extinction rates in Holarctic plants (Antonelli & Sanmartín, 2011; Willis & MacDonald, 2011; Xing et al., 2014). Willis & McDonald (2011) found that persistence and adaptation was the predominant response in the plant fossil record under Cenozoic climate warming. Here, we have found a mixed response of different clusioid lineages facing climate cooling, with evidence of extinction and/or migration (ancestors of the CPH clusioid lineages) but also of adaptation (*Hypericum* ancestors). This adaptation was probably crucial for the evolutionary success of *Hypericum*, the most species-rich and widespread of all clusioid genera. Yet, *Hypericum* might not be unique. Based on the Holarctic early Cenozoic fossil record, it has been argued that the climatic tolerances of many characteristic modern temperate taxa, including *Alnus, Quercus, Laurus, Fagus, Betula, Platanus, Carya, Ulmus* or *Vitis*, in North America, or *Populus, Salix, Myrica, Carpinus, Rubus, Ilex, Acer* or *Tilia* in Eurasia, changed in the Holarctic concurrent with the TEE (Wolfe, 1977). Further studies might shed light on whether these now temperate forest genera descend from ancient tropical taxa, which, as *Hypericum*, evolved temperate tolerances in *situ*.

To clarify this hypothesis for *Hypericum* and other such lineages, a combination of paleontological and neontological data might be crucial (Slater et al., 2012; Meseguer et al., 2015). If climate change is rapid and intense, it can have a clade-wide effect, driving to extinction entire lineages or promoting niche evolution across all members of a clade (Eiserhardt, Borchsenius, Plum, Ordonez & Svenning, 2015; Spriggs et al., 2015), in which case, evidence of ancestral climatic tolerances might be lost from extant taxa. In our reconstruction, where fossil information was limited to crown-node *Hypericum*, including evidence from extinct lineages did not change the inferred trend but contributed to decrease uncertainty in the inference (**Fig. 2**, **S4.3**). On the other hand, inclusion of fossil evidence might be critical to reconstruct the evolutionary fate of clades of the ancient, long-vanished "boreotropical" forest, such as the ancestors of the CPH clusioid clade, where extinction was inferred high.

## ACKNOWLEDGEMENTS

Research funding was provided by the Spanish government through a PhD grant (AP-2007-01698) to ASM, and projects CGL2009-13322-C01, CGL2012-40129-C01, and CGL2015-67849-P (MINECO/FEDER) to I.S. ASM was also supported by a Marie-Curie FP7-COFUND (AgreenSkills fellowship-26719). JC is supported by Marie-Curie actions People H2020, MSCA-IF-EF-ST-708207. The authors are grateful to Martin Godefroid, Graham Slater and to Sebastian Höhna for help with the analyses.

## LIST OF BRIEF TITLES OF ITEMS IN THE SI

Appendix S1: Clusioid CPH lineages occurrence dataset.

Appendix S2: Fossil *Hypericum* occurrence dataset.

Appendix S3: Extended Methods and Discussion.

Appendix S4: Supplementary Figures 1-7.

Appendix S5: Supplementary Tables 1-6.

Appendix S6: R script for diversification analyses in the package TESS.

Appendices S7-S10: for each variable, climatic values inferred using fossil calibrations.

Appendices S11-S14: for each variable, climatic values inferred without fossil calibrations.

## DATA ACCESIBILITY

All data used in this manuscript are presented in the manuscript and its supplementary material or have been previously published or archived elsewhere.

## BIOSKETCH

Andrea S. Meseguer is a postdoctoral researcher at ISEM (CNRS, France) interested in macroevolution, in the factors and processes generating geographic and diversity patterns. Isabel Sanmartín is a senior researcher at the Real Jardín Botánico (CSIC, Spain). Her main research interests include the study of large-scale biogeographical patterns, and the development of parametric methods in biogeography and macroevolution.

## AUTHOR CONTRIBUTIONS

ASM, JML and IS designed the study; DB provided paleoclimate data, and BRR and CCD the clusioid chronograms; ASM, JML, JC, and IS analysed the data; ASM and IS wrote the manuscript with contributions from JML, JC and EJ, and comments from all other authors.

## REFERENCES

Antonelli, A. & Sanmartín, I. (2011) Mass extinction, gradual cooling, or rapid radiation? Reconstructing the spatiotemporal evolution of the ancient angiosperm genus Hedyosmum (Chloranthaceae) using empirical and simulated approaches. Syst Biol, 60, 596–615.

Beerling, D., Berner, R.A., Mackenzie, F.T., Harfoot, M.B. & Pyle, J.A. (2009) Methane and the CH4 related greenhouse effect over the past 400 million years. Amer J Sci, 309, 97113.

Beerling, D.J., Fox, A., Stevenson, D.S. & Valdes, P.J. (2011) Enhanced chemistry-climate feedbacks in past greenhouse worlds. Proc Natl Acad Sci USA, 108, 9770–9775.

Betancur-R, R., Ortí, G. & Pyron, R.A. (2015) Fossil-based comparative analyses reveal ancient marine ancestry erased by extinction in ray-finned fishes. Ecol Lett, 18, 441–450.

Cavagnetto, C. & Anadón, P. (1996) Preliminary palynological data on floristic and climatic changes during the Middle Eocene-Early Oligocene of the eastern Ebro Basin, northeast Spain. Rev Palaeobot Palynol, 92, 281–305.

Couvreur, T.L., Pirie, M.D., Chatrou, L.W., Saunders, R.M., Su, Y.C., Richardson, J.E. & Erkens, R.H. (2011) Early evolutionary history of the flowering plant family Annonaceae: steady diversification and boreotropical geodispersal. J Biogeogr, 38, 664–680.

Crepet, W.L. & Nixon, K.C. (1998) Fossil Clusiaceae from the late Cretaceous (Turonian) of New Jersey and implications regarding the history of bee pollination. Am J Bot, 85, 1122–1133.

Davis, C.C., Bell, C.D., Mathews, S. & Donoghue, M.J. (2002) Laurasian migration explains Gondwanan disjunctions: Evidence from Malpighiaceae. Proc Natl Acad Sci U S A, 99, 6833–6837.

Davis, C.C., Webb, C.O., Wurdack, K.J., Jaramillo, C.A. & Donoghue, M.J. (2005) Explosive radiation of malpighiales supports a mid-Cretaceous origin of modern tropical rain forests. Am Nat, 165, E36–65.

Dawson, T.P., Jackson, S.T., House, J.I., Prentice, I.C. & Mace, G.M. (2011) Beyond predictions: biodiversity conservation in a changing climate. Science, 332, 53.

Donoghue, M.J. & Edwards, E.J. (2014) Biome shifts and niche evolution in plants. Annu Rev Ecol Evol Syst, 45, 547–542.

Dormann, C.F., Elith, J., Bacher, S., Buchmann, C., Carl, G., Carré, G., Marquéz, J.R.G., Gruber, B., Lafourcade, B., Leitão, P.J., Münkemüller, T., McClean, C., Osborne, P.E., Reineking, B., Schröder, B., Skidmore, A.K., Zurell, D. & Lautenbach, S. (2013) Collinearity: a review of methods to deal with it and a simulation study evaluating their performance. Ecography, 36, 27–46.

Eiserhardt, W.L., Borchsenius, F., Plum, C.M., Ordonez, A. & Svenning, J.-C. (2015) Climate-driven extinctions shape the phylogenetic structure of temperate tree floras. Ecol Lett, 18, 263–272.

Evans, M.E.K., Smith, S.A., Flynn, R.S. & Donoghue, M.J. (2009) Climate, niche evolution, and diversification of the “bird-cage” evening primroses (Oenothera, sections Anogra and Kleinia). Am Nat, 173, 225–240.

Graham, C.H., Carnaval, A.C., Cadena, C.D., Zamudio, K.R., Roberts, T.E., Parra, J.L., McCain, C.M., Bowie, R.C.K., Moritz, C., Baines, S.B., Schneider, C.J., VanDerWal, J., Rahbek, C., Kozak, K.H. & Sanders, N.J. (2014) The origin and maintenance of montane diversity: integrating evolutionary and ecological processes. Ecography, 37, 711–719.

Hewitt, G. (2000) The genetic legacy of the quaternary ice ages. Nature, 405, 907–913.

Hijmans, R.J., Cameron, S.E., Parra, J.L., Jones, P.G. & Jarvis, A. (2005) Very high resolution interpolated climate surfaces for global land areas. Int J Climatol, 25, 1965–1978.

Ho, L.S.T. & Ané, C. (2014) Intrinsic inference difficulties for trait evolution with Ornstein-Uhlenbeck models. Methods Ecol Evol, 5, 1133–1146.

Höhna, S., May, M.R. & Moore, B.R. (2016) TESS: an R package for efficiently simulating phylogenetic trees and performing Bayesian inference of lineage diversification rates. Bioinformatics, 32, 789–791.

Höhna, S., Landis, M.J., Heath, T.A., Boussau, B., Lartillot, N., Moore, B.R., Huelsenbeck, J.P. & Ronquist, F. (2016) RevBayes: Bayesian phylogenetic inference using graphical models and an interactive model-specification language. Syst Biol, 65, 726–736.

Lavin, M. & Luckow, M. (1993) Origins and relationships of tropical North America in the context of the boreotropics hypothesis. Am J Bot, 80, 1–14.

May, M.R., Höhna, S. & Moore, B.R. (2016) A Bayesian approach for detecting the impact of mass-extinction events on molecular phylogenies when rates of lineage diversification may vary. Methods Ecol Evol, 7, 947–959.

Merckx, V.S.F.T., Hendriks, K.P., Beentjes, K.K., Mennes, C.B., Becking, L.E., Peijnenburg, K.T.C.A., Afendy, A., Arumugam, N., de Boer, H., Biun, A., Buang, M.M., Chen, P.-P., Chung, A.Y.C., Dow, R., Feijen, F.A.A., Feijen, H., Soest, C.F.-v., Geml, J., Geurts, R., Gravendeel, B., Hovenkamp, P., Imbun, P., Ipor, I., Janssens, S.B., Jocque, M., Kappes, H., Khoo, E., Koomen, P., Lens, F., Majapun, R.J., Morgado, L.N., Neupane, S., Nieser, N., Pereira, J.T., Rahman, H., Sabran, S., Sawang, A., Schwallier, R.M., Shim, P.-S., Smit, H., Sol, N., Spait, M., Stech, M., Stokvis, F., Sugau, J.B., Suleiman, M., Sumail, S., Thomas, D.C., van Tol, J., Tuh, F.Y.Y., Yahya, B.E., Nais, J., Repin, R., Lakim, M. & Schilthuizen, M. (2015) Evolution of endemism on a young tropical mountain. Nature, 524, 347–350.

Meseguer, A.S. & Sanmartín, I. (2012) Paleobiology of the genus Hypericum (Hypericaceae): a survey of the fossil record and its palaeogeographic implications. An Jard Bot Madr, 69, 97–106.

Meseguer, A.S., Aldasoro, J.J. & Sanmartin, I. (2013) Bayesian inference of phylogeny, morphology and range evolution reveals a complex evolutionary history in St. John's wort (Hypericum). Mol Phylogenet Evol, 67, 379–403.

Meseguer, A.S., Lobo, J.M., Ree, R.H., Beerling, D.J. & Sanmartin, I. (2015) Integrating fossils, phylogenies, and niche models into biogeography to reveal ancient evolutionary history: the case of Hypericum (Hypericaceae). Syst Biol, 64, 215–232.

Morley, R.J. (2000) Origin and evolution of tropical rain forests. Wiley New York.

Morley, R.J. (2007) Cretaceous and Tertiary climate change and the past distribution of megathermal rainforests. Tropical Rainforest Responses to Climatic Change (ed. by M.B. Bush and J.-R. Flenley), pp. 1–31. Springer, Berlin.

Nürk, N.M., Uribe-Convers, S., Gehrke, B., Tank, D.C. & Blattner, F.R. (2015) Oligocene niche shift, Miocene diversification-cold tolerance and accelerated speciation rates in the St. John's worts (Hypericum, Hypericaceae). BMC Evol Biol, 15, 80–93.

O'Meara, B.C. (2012) Evolutionary inferences from phylogenies: a review of methods. Annu Rev Ecol Evol Syst, 43, 267–285

Otto-Bliesner, B.L., Marshall, S.J., Overpeck, J.T., Miller, G.H. & Hu, A. (2006) Simulating Arctic climate warmth and icefield retreat in the last interglaciation. Science, 311, 1751.

Paradis, E., Claude, J. & Strimmer, K. (2004) APE: Analyses of phylogenetics and evolution in R language. Bioinformatics, 20, 289–290.

Plummer, M., Best, N., Cowles, K. & Vines, K. (2006) CODA: convergence diagnosis and output analysis for MCMC. R News, 1, 7–11.

Pokorny, L., Riina, R., Mairal, M., Meseguer, A.S., Culshaw, V., Cendoya, J., Serrano, M., Carbajal, R., Ortiz, S., Heuertz, M. & Sanmartin, I. (2015) Living on the edge: timing of Rand flora disjunctions congruent with ongoing aridification in Africa. Front Genet, 6

Robson, N.K.B. (2012) Studies in the genus Hypericum L. (Hypericaceae) 9. Addenda, corrigenda, keys, lists and general discussion. Phytotaxa, 72, 1–111.

Ruhfel, B.R., Stevens, P.F. & Davis, C.C. (2013) Combined morphological and molecular phylogeny of the clusioid clade (Malpighiales) and the placement of the ancient rosid macrofossil Paleoclusia. Int J Plant Sci, 174, 910–936.

Ruhfel, B.R., Bove, C.P., Philbrick, C.T. & Davis, C.C. (2016) Dispersal largely explains the Gondwanan distribution of the ancient tropical clusioid plant clade. Am J Bot, 103, 1117–1128.

Ruhfel, B.R., Bittrich, V., Bove, C.P., Gustafsson, M.H.G., Philbrick, C.T., Rutishauser, R., Xi, Z. & Davis, C.C. (2011) Phylogeny of the clusioid clade (Malpighiales): Evidence from the plastid and mitochondrial genomes. Am J Bot, 98, 306–325.

Slater, G.J., Harmon, L.J. & Alfaro, M.E. (2012) Integrating fossils with molecular phylogenies improves inference of trait evolution. Evolution, 66, 3931–3944.

Spriggs, E.L., Clement, W.L., Sweeney, P.W., Madriñan, S., Edwards, E.J. & Donoghue, M.J. (2015) Temperate radiations and dying embers of a tropical past: the diversification of Viburnum. New Phytol, 207, 1469–8137.

Stevens, P.F. (2007) Clusiaceae-Guttiferae. The families and genera of vascular plants (ed. by K. Kubitzki). Springter, Berlin, Heidelberg.

Svenning, J.-C., Eiserhardt, W.L., Normand, S., Ordonez, A. & Sandel, B. (2015) The influence of paleoclimate on present-day patterns in biodiversity and ecosystems. Annu Rev Ecol Evol Syst, 46, 551–572.

Tiffney, B.H. (1984) Seed size, dispersal syndromes, and the rise of the angiosperms: Evidence and hypothesis. Ann Mo Bot Gard, 71, 551–576.

Tiffney, B.H. (1985) Perspectives on the origin of the floristic similarity between eastern Asia and eastern North America. J Arnold Arbor, 66, 73–94.

Töpel, M., Zizka, A., Calió, M.F., Scharn, R., Silvestro, D. & Antonelli, A. (2017) SpeciesGeoCoder: fast categorization of species occurrences for analyses of biodiversity, biogeography, ecology, and evolution. Syst Biol, 66, 145–151.

Upchurch, G.R. & Wolfe, J.A. (1987) Mid-Cretaceous to early Tertiary vegetation and climate: Evidence from fossil leaves and woods. The origins of Angiosperms and their biological consequences (ed. by E.M. Friis, W.G. Chaloner and J.C. Crane), pp. 75–105. Cambridge University Press, Cambridge.

Wiens, J.J., Ackerly, D.D., Allen, A.P., Anacker, B.L., Buckley, L.B., Cornell, H.V., Damschen, E.I., Jonathan Davies, T., Grytnes, J.-A., Harrison, S.P., Hawkins, B.A., Holt, R.D., McCain, C.M. & Stephens, P.R. (2010) Niche conservatism as an emerging principle in ecology and conservation biology. Ecol Lett, 13, 1310–1324.

Willis, K.J. & MacDonald, G.M. (2011) Long-term ecological records and their relevance to climate change predictions for a warmer world. Annu Rev Ecol Evol Syst, 42, 267287.

Wolfe, J.A. (1975) Some aspects of plant geography of the Northern Hemisphere during the Late Cretaceous and Tertiary. Ann Mo Bot Gard, 62, 264–279.

Wolfe, J.A. (1977) Paleogene floras from the Gulf of Alaska region. U.S. Geological Survey, 997, 1–108.

Wolfe, J.A. (1985) Distribution of major vegetational types during the Tertiary. In: The carbon cycle and atmospheric CO_2_: Natural variations archean to present, pp. 357–375. Proceedings of the Chapman Conference on Natural Variations in Carbon Dioxide and the Carbon Cycle, Tarpon Springs, FL.

Wolfe, J.A. & Upchurch, G.R. (1987) North American nonmarine climates and vegetation during the Late Cretaceous. Palaeogeogr Palaeoclimatol Palaeoecol, 61, 33–77.

Wüest, R.O., Antonelli, A., Zimmermann, N.E. & Linder, H.P. (2015) Available climate regimes drive niche diversification during range expansion. Am Nat, 185, 640–652.

Wurdack, K.J. & Davis, C.C. (2009) Malpighiales phylogenetics: Gaining ground on one of the most recalcitrant clades in the angiosperm tree of life. Am J Bot, 96, 1551–1570.

Xi, Z., Ruhfel, B.R., Schaefer, H., Amorim, A.M., Sugumaran, M., Wurdack, K.J., Endress, P.K., Matthews, M.L., Stevens, P.F., Mathews, S. & Davis, C.C. (2012) Phylogenomics and a posteriori data partitioning resolve the Cretaceous angiosperm radiation Malpighiales. Proc Natl Acad Sci USA, 109, 17519–17524.

Xing, Y., Onstein, R.E., Carter, R.J., Stadler, T. & Linder, P.H. (2014) Fossils and a large molecular phylogeny show that the evolution of species richness, generic diversity, and turnover rates are disconnected. Evolution, 68, 2821–2832.

Zachos, J.C., Dickens, G.R. & Zeebe, R.E. (2008) An early Cenozoic perspective on greenhouse warming and carbon-cycle dynamics. Nature, 451, 279–283.

Zanne, A.E., Tank, D.C., Cornwell, W.K., Eastman, J.M., Smith, S.A., FitzJohn, R.G., McGlinn, D.J., O/'Meara, B.C., Moles, A.T., Reich, P.B., Royer, D.L., Soltis, D.E., Stevens, P.F., Westoby, M., Wright, I.J., Aarssen, L., Bertin, R.I., Calaminus, A., Govaerts, R., Hemmings, F., Leishman, M.R., Oleksyn, J., Soltis, P.S., Swenson, N.G., Warman, L. & Beaulieu, J.M. (2014) Three keys to the radiation of angiosperms into freezing environments. Nature, 506, 89–92.

Ziegler, A., Eshel, G., Rees, P.M., Rothfus, T., Rowley, D. & Sunderlin, D. (2003) Tracing the tropics across land and sea: Permian to present. Lethaia, 36, 227–254.

